# Discrete and coordinated encoding of punishment contingent on rewarded actions by prefrontal cortex and VTA

**DOI:** 10.1101/157032

**Authors:** Junchol Park, Bita Moghaddam

## Abstract

Actions motivated by a rewarding outcome are often associated with a risk of punishment. Little is known about the neural representation of punishment that is contingent on reward-guided behavior. We modeled this circumstance by using a task where actions were consistently rewarded but probabilistically punished. Spike activity and local field potentials were recorded during this task simultaneously from VTA and mPFC, two reciprocally connected regions implicated in both reward-seeking and aversive behavioral states. At the single unit level, we found that ensembles of VTA and mPFC neurons encode the contingency between action and punishment. At the network level, we found that coherent theta oscillations synchronize the VTA and mPFC in a bottom-up direction, effectively phase-modulating the neuronal spike activity in the two regions during punishment-free actions. This synchrony declined as a function of punishment contingency, suggesting that during reward-seeking actions, risk of punishment diminishes VTA-driven neural synchrony between the two regions.

## Introduction

Goal-directed actions aimed at obtaining a reward often involve exposure to an aversive event or punishment. For example, foraging for food in the wild may result in encountering a predator. In a causally and socially complex world, appropriate representation of punishment that is contingent on reward-seeking actions is critical for survival and optimal action selection. Deficits in this representation may be associated with detrimental behavioral patterns observed in addictive disorders while exaggerated representation of punishment may be linked to anxiety-related disorders (Bechara et al., 2000; Gillan et al., 2016; Hartley and Phelps, 2012; Lee, 2013; Mineka et al., 1998).

How is punishment that is contingent on reward-seeking actions represented by the brain? To begin to address this question, we focused on the ventral tegmental area (VTA) and the medial prefrontal cortex (mPFC). Neurons in the VTA including dopamine (DA) and non–dopamine neurons are critical components of the reward circuitry including reward-mediated actions (Cohen et al., 2012; Matsumoto et al., 2016; Roesch et al., 2007; Schultz, 1998; Tan et al., 2012; van Zessen et al., 2012; Wise, 2004). We, therefore, hypothesized that ensembles of VTA neurons represent risk of punishment associated with reward-mediated behavior. Importance of VTA notwithstanding, the mPFC is also implicated in encoding of reward and reward-guided action selection (Barraclough et al., 2004; Buschman et al., 2012; Kobayashi et al., 2006; Powell and Redish, 2016; Rich and Shapiro, 2009), as well as control of aversive and anxiety-like behavior (Adhikari et al., 2010; Karalis et al., 2016b; Kumar et al., 2014; Likhtik et al., 2014; Park et al., 2016; Ye et al., 2016). At a circuit level, the mPFC and VTA (both DA and non-DA neurons) send reciprocal projections to each other (Berger et al., 1976; Carr and Sesack, 2000a, b). The VTA neurons projecting to mPFC have been shown to respond to stressful and anxiogenic perturbations with a greater degree of sensitivity compared to the mesolimbic or mesostriatal projections (Abercrombie et al., 1989; Bradberry et al., 1991; Moghaddam et al., 1990; Thierry et al., 1976). Furthermore, photostimulation of the VTA dopaminergic input to the mPFC elicits anxiety-like behavior (Gunaydin et al., 2014; Lammel et al., 2012), suggesting a more causal role for this neural circuit in aversive behavior. Given this, we further hypothesized that the interaction between VTA and mPFC provides a dynamic representation of punishment contingent on a reward-seeking action and punishment-based modulation of that action.

To test these hypotheses, we first designed and validated a task that allowed us to assess reward-directed instrumental behavior in the absence or presence of action-contingent punishment in the same recording session. The latter criterion was critical because it allowed us to track the activity of the same ensembles of neurons as the punishment contingency associated with the same instrumental behavior changed. The task was designed so that the action always procured a reward but the same action probabilistically led to punishment with block-wise varying degrees of contingency. Thus, different blocks had varying action-punishment contingency whereas the action-reward contingency remained constant. We then recorded single unit activity and local field potentials (LFP) from the VTA and mPFC simultaneously during this task. The simultaneous recording allowed us to characterize and compare the inter- and intra-regional neural codes for punishment and punishment-based modulation of instrumental action.

At the single unit level, we found that VTA and mPFC neurons encode action-punishment contingency and punishment-based behavioral modulation. At the network level, we found that coherent theta oscillations synchronize the VTA and mPFC in a bottom-up direction, effectively phase-modulating the neuronal spike activity in the two regions during punishmen-free actions. This oscillation-mediated neural synchrony declined as a function of action-punishment contingency, suggesting that desynchronization of the two regions signals punishment.

## Results

### Anxiety-like behavioral changes by punishment contingent on the action

The conditional probability of punishment contingent on the action – termed ‘punishment’ hereafter for simplicity – induced anxiety-like aversive behavioral changes (Figure 1a-b), as the mean response time (RT) markedly increased as a function of punishment (Figure 1c; GLM repeated measures, *F*_2,32_ = 24.94, *p* < 0.001). We also measured time spent in immobility during RT, a widely used behavioral measure of anxiety in rodents, and observed a significant increase in immobile RT (Figure 1c; GLM repeated measures, *F*_2,32_= 22.44, *p* < 0.001),demonstrating that the increased RT may involve anxiety. The increase in RT could not be explained by changes in motivation for the reward, as the time for reward retrieval (reward RT) remained consistent across blocks regardless of punishment (Figure 1c inset; GLM repeated measures, *F* _2,32_ = 2.97, p = 0.07). When animals performed the same number of trials and blocks in the absence of punishment throughout a session, namely ‘no-shock control session’, the mean RT and reward RT did not differ across blocks (Figure 1d).

**Figure 1.**
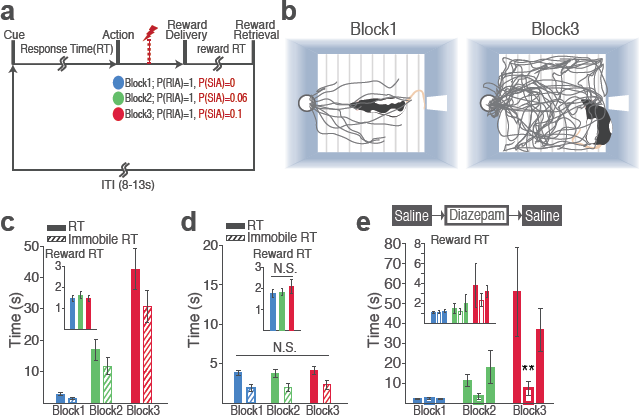
Punishment induces anxiety-like changes in reward-seeking behavior. (a) A schematic diagram illustrating the task. (b) Representative behavioral trajectories in block 1 (left, 10 trials) and block 3 (right, 10 trials). (c) Significant increases in RT (filled bars) and immobile RT (slashed bars) were observed as a function of punishment contingency. (Inset) Latency from reward delivery to retrieval (reward RT) did not differ across blocks. (d) RT, immobile RT, and reward RT did not change across blocks in the absence of punishment. (e) Animals performed three sessions of the task with pretreatment of saline (Day 1) - diazepam (2 mg/kg) - saline (Day 2). Pretreatment of an anxiolytic diazepam (2 mg/kg) but not saline injection averted punishment-induced increase in the mean RT. **p < 0.005; post hoc test. (Inset) Injections did not influence reward RT.

The punishment-induced increase in RT was pertinent to anxiety because an anxiolytic drug, diazepam, reduced the increase in RT. Systemic pretreatment of diazepam (2 mg/kg) significantly averted increase in RT of block 3, compared with saline-pretreatment data (Figure 1e; Repeated measures ANOVA, *F*_4,48_ = 3.27, *p* = 0.019; *post hoc test,* block 3, *p* values < 0.01). Diazepam or saline injected animals showed equivalent levels of reward RT (Figure 1e inset; Repeated measures ANOVA, *F*_4,48_ = 0.34, *p* = 0.852; *post hoc test, p* values > 0.51).

### Individual neuronal encoding of punishment

During task performance, 167 mPFC and 102 VTA single units were recorded from histologically verified electrodes (Figure S1). For all single unit data analyses, we classified VTA units into putative dopamine (DA, n = 55) and putative non-dopamine (non-DA, n = 47) subtypes (Figure S2, Experimental procedure). We first examined the trial-averaged neuronal activity of mPFC, VTA DA, and non-DA units to compare their general tuning properties during task events – cue onset, action, and reward delivery. Figure 2a shows peri-event neuronal activity averaged across all trials and blocks. The majority of VTA DA units displayed phasic excitatory responses at each task event as has been previously reported (Schultz et al., 1993), whereas non-DA and mPFC units showed weaker and temporally diffuse responses (Figure 2a-c; Repeated measures ANOVA, post-cue, *F*_2,226_ = 22.05; peri-action, *F*_2,266_ = 43.78; post-reward, *F*_2,266_ = 48.93, *p* values < 0.001).

**Figure 2.**
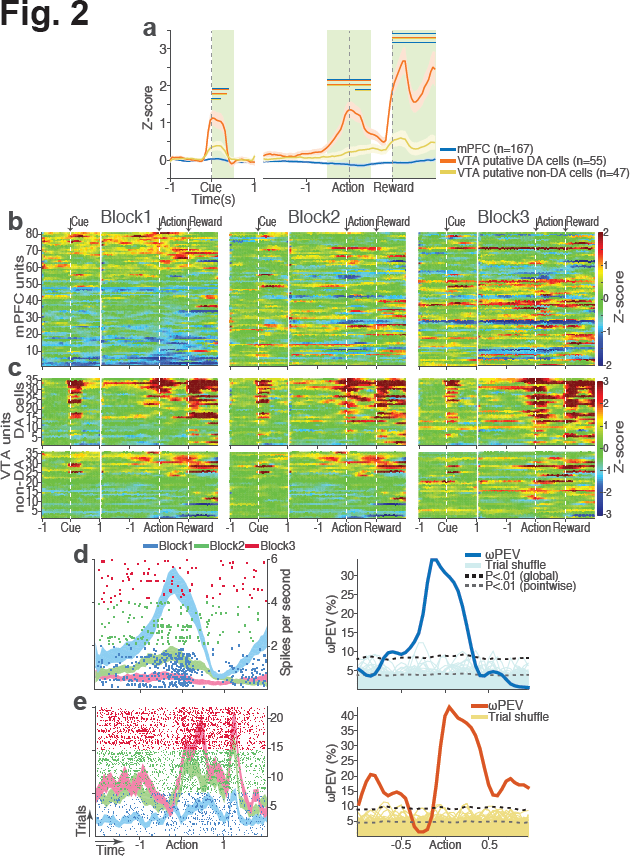
mPFC, VTA DA, and non-DA single units respond to task events and punishment. (a) Peri-event activity averaged across all trials and all units within each neuron group. Dual–colored bars above indicate significant pairwise differences at corresponding time bins according to the post hoc analysis (p < 0.05). The green shadows indicate time windows of statistical analyses. (b) Baseline-normalized peri-event firing rates of mPFC units are plotted per block to reveal neuronal responses to punishment. Only units with significant activity modulation are plotted - i.e., punishment-encoding units (Figure S3). (c) Peri-event activity of VTA putative DA (top panels) and non-DA (bottom panels) punishment-encoding units. (d-e) Identification of single units discriminating their firing rates across different blocks as a function of punishment.(d)Left, A raster plot showing a representative mPFC unit’s peri-action spike activity across blocks with spike density functions of different blocks superimposed. Right, To quantify each unit’s encoding, percent variance in the unit’s firing rate explained by blockwise change in punishment contingency (wPEV) was calculated. To determine the global wPEV band, trial-shuffled surrogate wPEV distribution (light blue curves) was acquired, and the pointwise and global wPEV bands were found from the distribution at a = 0.01 (Experimental procedure). A unit whose ωPEV curve crosses the global band was determined as a punishment-encoding unit. (e) Left, A representative VTA unit’s peri-action activity across blocks. Right, This VTA unit satisfied the punishment-encoding criterion.

We then examined modulation of single neuronal activity across blocks (Figure 2b-c). Some of mPFC, VTA DA, and non-DA single units modulated their peri-event firing rates across blocks as a function of action-punishment contingency. We therefore quantified individual neuronal encoding of punishment using a percent explained variance (ωPEV) statistic (Buschman et al., 2012) - i.e., how much of total variance in a neuron’s firing rate across trials can be explained by punishment contingency varying across blocks (Experimental procedure). To identify punishment-encoding units, we compared the ωPEV statistic of the original spike trains with the ωPEV distribution of surrogate spike trains created by shuffling block labels (Figure 2d-e, Experimental procedure). Substantial proportions of units in both regions (mPFC, 49 %; VTA DA, 65 %; VTA non-DA, 77 %) encoded punishment at the time of the action (Figure 3a, S3). Likewise, units displayed more pronounced encoding of punishment at the time of the action, since the peri-action ωPEVs appeared to be greater than that of the peri-cue or peri-reward epoch (Figure 3a). Therefore, subsequent analyses focused on the peri-action epoch.

**Figure 3.**
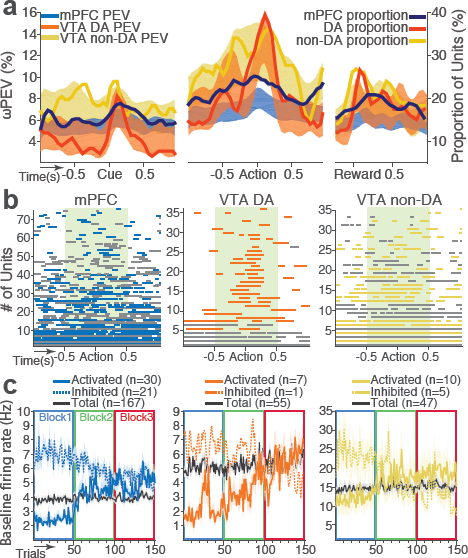
mPFC, VTA DA, and non-DA single units encode action-punishment contingency by modulating their peri-event and baseline firing rates. (a) Shaded area indicates the mean ± s.e.m. ωPEV averaged across all units in each neuron group (left vertical axis) across time. Line plots indicating proportions of punishment-encoding units are superimposed (right vertical axis). (b) To reveal time points of punishment encoding, crossing of the global ωPEV band by each unit is marked with a line segment (Experimental procedure). Only the units with at least one crossing are included in each plot. Single units with significant change in their baseline firing rate are marked with gray lines (see below). (c) Subpopulations of single units represented punishment with significant excitatory or inhibitory modulation of their baseline (inter-trial interval) activity. The mean ± s.e.m. baseline firing rates are plotted across trials and blocks.

mPFC, VTA DA, and non-DA units exhibited equivalent levels of punishment encoding in the peri-action epoch given that the main effect of unit groups was not significant (*F*_2,266_ = 1.21, *p* = 0.30). However, a significant interaction between time and unit groups was observed in the peri-action ωPEV (Figure 3a; Repeated measures ANOVA, *F*_38 5054_ = 2.68, *p* < 0.001), indicating distinct time-varying patterns in encoding of punishment by different unit groups. We then examined the time course of individual neuronal encoding in the peri-action epoch. We recalculated the peri-action ωPEV using a narrower moving window (50-ms width, 5-ms step) to reveal time points of individual neuronal encoding of punishment with a higher degree of temporal precision. VTA DA neuronal encoding appeared to be concentrated specifically around the time of the action, whereas non-DA and mPFC units displayed temporally diffuse patterns (Figure 3b). This result suggests that DA units may be involved in more precise signaling of action-punishment contingency whereas the mPFC and VTA non-DA units may represent more persistent effects of punishment on motivational/emotional states. Consistent with this function, substantial proportions of mPFC (31 %) and non-DA (32 %) units significantly modulated their baseline firing rates across blocks, suggesting representation of punishment on a larger temporal scale (Figure 3c). Fewer VTA DA units (14 %) modulated their baseline activity (Figure 3c).

We examined the direction of activity modulation (excitation or inhibition) as a function of punishment. Similar proportions of mPFC units encoded punishment with bidirectional modulation of activity (Figure 4a), which resulted in a lack of net excitation or suppression of activity across blocks (Figure 4b; Repeated measures ANOVA, *F*_2,241_ = 2.09, *p* = 0.126). The majority of VTA DA units encoded punishment contingency with an excitatory modulation of activity (Figure 4c) specifically at the time of the action (Figure 4d; Repeated measures ANOVA, 2 105 = 3.96, *p* = 0.022), but not during other task events. Similar to the DA units, the greater number of non-DA punishment-encoding units displayed an excitatory modulation of activity (Figure 4e) with a trend toward a net excitatory effect of the block (Figure 4f; Repeated measures ANOVA, *F*_2,105_. = 2.94, *p* = 0.057).

**Figure 4.**
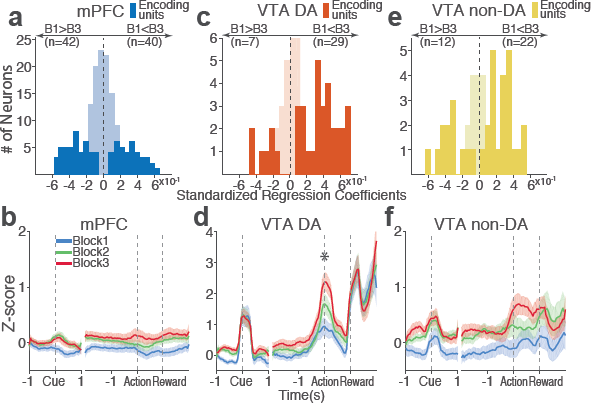
Distinct subpopulations of single units encode action-punishment contingency with excitatory or inhibitory activity modulation. (a, c, e) Units are distributed across the horizontal axis based on modulation of their peri-action activity across blocks as a function of punishment. Standardized regression coefficients (SRC) were computed for a normalized quantification of each unit’s peri-action activity modulation by punishment (Experimental procedure). In each distribution, units with excitatory or inhibitory activity modulation are located in the right or left portion of the distribution, respectively. Punishment-encoding units are solid-colored, while non-encoding units are pale-colored. (a) Direction of the mPFC neuronal activity modulation. (b) The baseline-normalized activity of the mPFC encoding units per block (mean ± s.e.m.). (c) Direction of the VTA DA neuronal activity modulation. (d) The activity of the VTA DA encoding units per block (mean ± s.e.m.). Asterisk indicates a significant effect of block on the peri-action activity (p <0.05). (e) Direction of the VTA non-DA neuronal activity modulation. (f) The activity of the VTA non-DA encoding units per block (mean ± s.e.m.).

To disentangle neuronal encoding of punishment from other confounding factors that may cause block-dependent changes in neuronal activity – such as satiety, fatigue, or spontaneous changes over time, we performed a control experiment recording from 126 mPFC and 57 VTA single units (putative DA, n = 28; putative non-DA, n = 29) during performance of three consecutive punishment-free blocks. We observed a much weaker blockwise changes in firing rate indicating negligible impact of confounding factors on the neuronal encoding of punishment (Figure 5).

**Figure 5.**
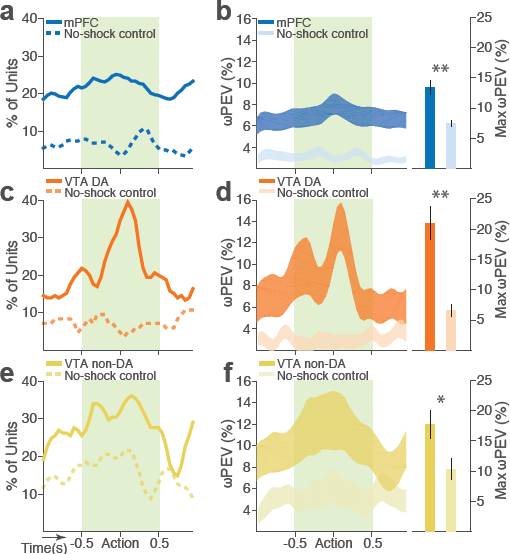
Blockwise firing rate changes in the presence vs absence of punishment contingency. (a)Proportion of mPFC units showing significant firing-rate changes across blocks during the peri-action epoch in the presence vs absence (no-shock control) of punishment. (b) Left,Percent variance in the mPFC unit firing rates explained by the block shift (ωPEV) in the presence vs absence of punishment (mean ± s.e.m.). Right, Maximum peri-action ωPEV of mPFC units differed in the presence vs absence of punishment (Student’s t-test, t = 3.81, *p* < 0.001). (c) Proportion of VTA DA units showing significant firing-rate changes across blocks. (d) Left, ωPEV of VTA DA units. Right, Maximum peri-action ωPEV of VTA DA units (t_81_ = 4.19, *p* < 0.001). (e) Proportion of VTA non-DA units showing significant firing-rate changes across blocks. (f) Left, ωPEV of VTA non-DA units. Right, Maximum peri-action ωPEV of VTA non-DA units (t_74_ = 2.25, *p* = 0.028). *p < 0.05, **p < 0.005.

Taken together, these data indicate that both VTA and mPFC single neurons convey the information about action-punishment contingency. Moreover, we observed distinct temporal and directional tuning properties from mPFC, VTA DA, and non-DA neuronal subpopulations. This heterogeneity may enable the VTA-mPFC circuit to represent diverse motivational/emotional aspects of punishment.

### Contingency of punishment diminishes neural synchrony between VTA and mPFC

At the neural circuit and network level, synchronous oscillations can provide temporal coordination among local and interregional neural groups, thereby promoting various cognitive, emotional, and motivational functions (Adhikari et al., 2010; Buschman et al., 2012; Fries, 2015; Karalis et al., 2016a; Kim et al., 2012; Likhtik et al., 2014). We examined whether such oscillation-mediated neural synchrony subserved encoding of reward-seeking action and punishment contingent on the action.

During punishment-free performance in block 1, theta oscillations in the frequency band of 5 to 15 Hz emerged in mPFC and VTA both preceding and during the action (Figure 6a-d). This theta oscillation was markedly reduced as a function of punishment contingency in both regions (Figure 6e-f). Of note, this change in theta oscillation observed before and during the action cannot be explained by the discrepancy in the animals’ motor activity because they engaged in the action within this epoch. A significant interaction between punishment and frequency band was detected in LFP power during post-cue and pre-action time periods in both regions (Repeated measures ANOVA, VTA, post-cue, *F*_52, 1092_ = 5.1 1; pre-action, *F*_52, 1092_ = 7.8, *p* values < 0.001; mPFC, post-cue, *F*_52 1872_ = 1.60, *p* = 0.005; pre-action, *F*_52 1872_ = 2.29, *p* <0.001, Figure 6g-h). Theta oscillations appeared to be coherent between the two regions during the punishment-free action, but the coherence significantly reduced as a function of punishment (Figure 6i). A significant interaction between punishment and frequency band was observed in LFP coherence during the pre-action period (Repeated measures ANOVA, post-cue, *F*_52 1092_ = 0.93, *p* = 0.62; pre-action, *F*_52 1092_ = 2.45, *p* < 0.001).

**Figure 6.**
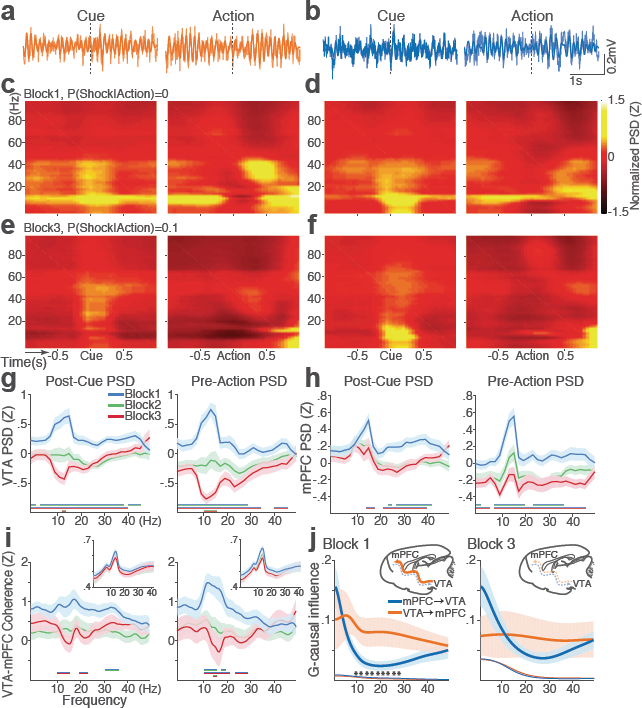
Punishment diminishes theta oscillation-mediated neural synchrony in the VTA-mPFC circuit. (a) Representative VTA peri-event LFP traces in a block 1 trial. Bandpass filtered LFP signal (heavy line) is superimposed on the raw trace (thin line). (b) Simultaneously recorded mPFC LFP traces. (c) Baseline-normalized VTA power spectrograms averaged across block 1 trials (left: peri-cue, right: pre-action). mPFC block 1 power spectrograms are in (d). (e) Diminished VTA theta power in block 3. (f) Similar diminishment observed in mPFC theta power. (g) Mean ± s.e.m. (shaded area) normalized VTA PSDs per block corresponding to 1-s post-cue (left) and pre-action (right) epochs. Dual-colored bars below indicate significant pairwise differences at corresponding frequency bins according to post hoc analyses (p < 0.05). (h) Normalized mPFC PSDs in post-cue (left) and pre-action (right) epochs. (i) Normalized VTA-mPFC LFP coherence in post-cue (left) and pre-action (right) epochs. Insets represent non–normalized LFP coherences of each block. (j) Granger-causality, representing mutual influences (directionality) between VTA and mPFC peri-action LFP time series in block 1 (left) and block 3 (right). Blue and orange curves represent mPFC-to-VTA and VTA-to-mPFC Granger-causal influences, respectively. Shaded areas indicate s.e.m. Thin colored-lines below indicate upper bounds of confidence intervals (a = 0.001) acquired by the random permutation resampling of time bins. Asterisk indicates significant difference between bidirectional Granger-causal influences at the corresponding frequency bin (p < 0.05).

To examine mutual influences (directionality) of LFP time series between the two regions, we quantified Granger causal influences (GC) in VTA-to-mPFC and mPFC-to-VTA directions (Experimental procedure). During punishment-free action in block 1, the theta oscillation was driven by VTA, as mPFC was GC influenced by VTA significantly greater than the GC influence of the other direction (Figure 6j). A significant interaction between directionality and frequency band was observed in GC coefficients in all blocks, indicating that the oscillation directionality varied across frequency bands (Figure 6j; Repeated measures ANOVA, Block 1, *F*_25,700_ = 2.05, *p* = 0.002; Block 2, *F*_25,700_ = 2.38, *p* < 0.001; Block 3, *F*_25,700_ = 5.59, *p* < 0.001).But importantly post hoc analysis revealed significantly greater VTA-to-mPFC directionality in frequency bands including the theta band in block 1, and the directionality became unclear in blocks 2 & 3 (Figure 6j, data not shown for block 2). Taken together, these results suggest that the VTA-driven theta oscillation entrains the VTA-mPFC circuit during punishment-free action. Decline in this entrainment may represent punishment contingent on the action, since power, coherence, and directionality of the theta oscillation declined as a function of punishment contingency. Analyses of no-shock control data revealed that these punishment-dependent changes in the VTA-mPFC theta oscillations were not evident in the absence of punishment (Figure S4).

### Punishment-induced reduction in local and interregional LFP–spike synchrony

Synchronous oscillations can provide temporal coordination of spike activity of local and interregional neuron groups by creating rhythmic sequences of neuronal excitation and inhibition, thereby enhancing ‘neuronal communication’ between coherently timed neuron groups (Fries, 2015; Harris and Gordon, 2015). The presence of such LFP-mediated spike timing coordination was examined by measuring phase-locking of the neuronal spike activity to local and interregional theta oscillations in the peri-action epoch.

Within each region, substantial proportions of single units (37 % in VTA and 23 % in mPFC) that were subjected to the phase-locking analysis showed significant phase-locking to the local theta oscillation in the punishment-free block 1 according to Rayleigh’s z-test for non–uniformity of the spike phase distribution. Consistent with the temporally specific increase in the theta spectral power (Figure 6c-d), the phase synchrony arose during action from the baseline level in the phase-locked units in both regions (Figure 7a & d; Signed-rank test, *p* values < 0.001). Enhanced phase-locking, albeit to a lesser degree, was observed even in units that failed the Rayleigh’s test (Figure 7a & d; Signed-rank test, *p* values < 0.001), indicating widespread influence of the theta oscillation on local neuronal spike timing. The temporal relationship (directionality) between the spike outputs and the theta oscillation was examined using a time-lagged phase-locking analysis (Likhtik et al., 2014; Spellman et al., 2015). The spike-LFP phase-locking was recalculated using spike times shifted relative to the local or interregional LFP to infer the directionality in the LFP-spike interaction. We found that in block 1 greater proportions of units appeared to be phase-locked with negative time lags. The vast majority of phase-locked units (VTA, 74 %; mPFC, 67 %) had their maximum phase-locking values (PLVs; Experimental procedure) with a negative lag (Figure 7b & e; Signed-rank test, *p*values < 0.005). These indicated an entrainment of spike timing by preceding cycles of the oscillation - i.e., directionality from the theta oscillation to the spike activity. To examine the modulation of LFP-spike phase-locking by punishment, we compared PLVs across different blocks. A trend toward reduction in PLV was found in block 3 compared with block 1 in mPFC (Figure 7c; Signed-rank test, *p* = 0.077), and a significant reduction was found in VTA (Figure 7f; *p* = 0.006). Likewise, a trend toward reduction in the proportion of phase-locked units was observed in block 3 compared with block 1 in mPFC (Figure 7b; Chi-square test, y^2^ = 3.25, *p* = 0.071), and a significant reduction was found in VTA (Figure 7e; y^2^ = 4.31, *p* = 0.038). We next examined VTA DA and non-DA neuronal phase-locking separately. Greater fraction of DA units (45 %) appeared to be phase-locked compared with non-DA units (23 %) in block 1 (Figure 7g; Chi-square test, X^2^_1_ = 5.04, *p* = 0.025). The DA neuronal PLV in block1 significantly declined as a function of punishment contingency in block 2 & 3 (Figure 7h; Signed-rank test, *p* values < 0.01), whereas the non-DA neuronal PLV did not differ across blocks (p values > 0.43). These indicated that the punishment-induced reduction in the VTA neuronal phase-locking was predominately due to the reduction in DA neuronal synchrony.

**Figure 7.**
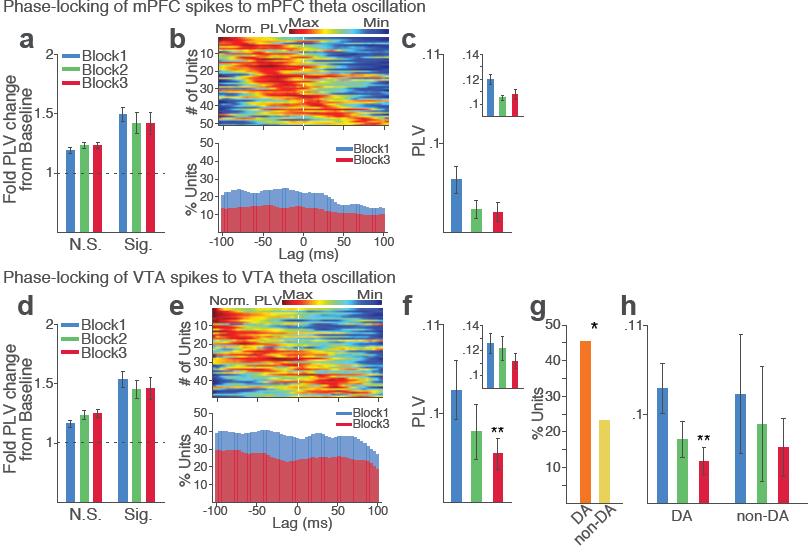
Contingency of punishment reduces VTA and mPFC neuronal synchrony to local theta oscillation. (a-c) Modulation of mPFC neuronal synchrony to mPFC theta oscillation. Phase-locking values (PLVs) were quantified by averaging 1,000 mean resultant lengths (MRLs) of the circular phase angle distribution comprising 100 resampled spikes per iteration (Experimental procedure). (a) Fold change from baseline in the strength of the neuronal phase-locking during peri-action epoch in units that passed Rayleigh z-test (Sig.) and rest of the units (N.S.). (b) Top, Normalized PLVs in block 1 across a range of time lags for all phase-locked mPFC units, aligned by peak lags. Bottom, Percentage of significantly phase-locked mPFC units in block 1 vs 3 across a range of lags. (c) Mean ± s.e.m. PLVs across different blocks. Inset, PLVs including significantly phase-locked units only. (d-h) Modulation of VTA neuronal synchrony to VTA theta oscillation. (d) Fold change from baseline in the strength of the neuronal phase-locking. (e) Top, Normalized PLVs in block 1 of all phase-locked VTA units. Bottom, Percentage of significantly phase-locked VTA units. (f) Mean ± s.e.m. PLVs across different blocks. (g) Percentage of phase-locked VTA DA and non-DA units. (h) PLVs of VTA DA and non-DA units plotted separately.

Next we examined the interregional LFP-spike phase-locking between VTA and mPFC. Based on the Granger causal influence indicating VTA-to-mPFC directionality in theta oscillations, we anticipated stronger mPFC neuronal synchrony to the VTA theta oscillation than that of the other direction. Consistent with this, we found that a substantial proportion of mPFC units (31 %) were phase-locked to the VTA theta oscillation in block 1. A representative mPFC unit with significant phase-locking is shown in Figure 8a-b. The interregional spike-phase synchrony emerged during the action compared to the baseline (Figure 8c, Signed-rank test, *p*values < 0.001). We examined directionality of the LFP-spike synchrony using the time-lagged phase-locking analysis. In block 1, the majority of phase-locked units had their peak PLVs with a negative lag (Figure 8d; Signed-rank test, *p* = 0.066). Likewise, greater proportions of phase-locked units were observed on negative lags (Figure 8d, bottom). In addition, the mean PLV across negative time lags appeared to be greater than that of the positive lags (Figure 8e; Signed-rank test, *p* = 0.023). These indicate mPFC neuronal entrainment to preceding VTA theta oscillatory cycles – i.e., VTA-to-mPFC directionality. When compared across blocks, the mPFC neuronal entrainment by the VTA theta oscillation declined as a function of punishment contingency (Figure 8e-g, Signed-rank test, *p* = 0.003). As the degree of phase-locking diminished, the VTA-to-mPFC directionality also declined (Figure 8d-e). We also examined the VTA neuronal phase-locking to the mPFC theta oscillation. The degree of VTA neuronal phase-locking to the mPFC theta oscillation appeared to be much weaker than the mPFC neuronal phase-locking to the VTA theta oscillation (Wilcoxon Rank-sum test, *p* < 0.001), corroborating the VTA-to-mPFC directionality in the theta-oscillation-mediated spike phase modulation (Figure S5). The PLVs did not differ across different blocks in both DA and non-DA units (Figure S5). Analyses of no-shock control data showed unchanging neural synchrony across blocks in the absence of punishment (Figure S6-7).

**Figure 8.**
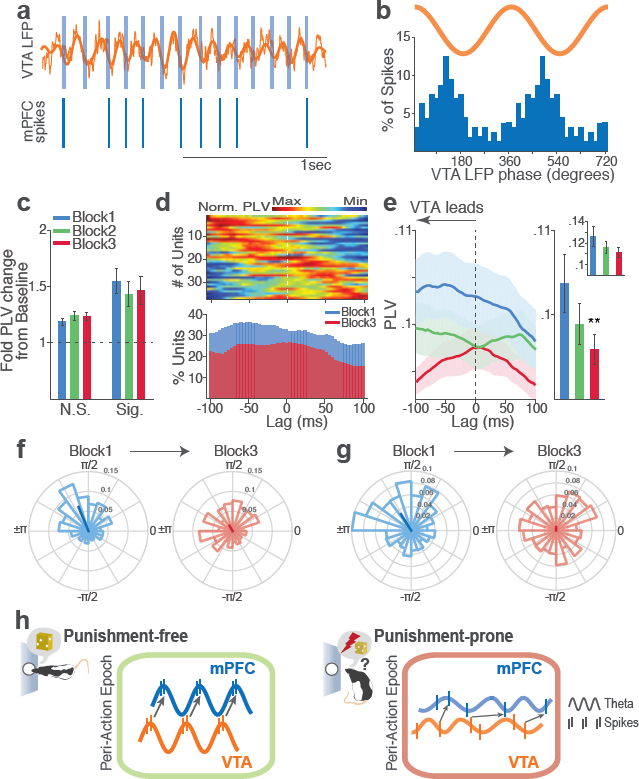
Contingency of punishment reduces mPFC neuronal synchrony to the VTA theta oscillation. (a) Top, Example raw (thin line) and bandpass filtered (heavy line) VTA LFP traces. Bottom, Neuronal spikes of a simultaneously recorded mPFC single unit. This unit’s preferred phase is indicated with light blue columns superimposed on the LFP trace. (b) Distribution of spike phase angles of the example mPFC unit relative to the VTA theta oscillation (Rayleigh’s *p* <0.001). (c) Fold change from baseline in the strength of mPFC neuronal phase-locking (PLV) during the peri-action epoch in units that passed Rayleigh z-test (Sig.) and rest of the units (N.S.). (d) Top, Normalized PLVs in block 1 across a range of time lags for all phase-locked mPFC units, aligned by peak lags. Bottom, Percentage of significantly phase-locked units in block 1 vs 3 (e) Left, PLVs calculated with negative and positive time lags applied to spike trains relative to LFP time series. Right, Mean ± s.e.m. PLVs of all units across different blocks. Inset, PLVs including significantly phase-locked units only. (f-g) Polar plots represent the distribution of spike-phase angles of an example mPFC unit in block 1 vs 3. To quantify the circular concentration of phase angles, we calculated the mean resultant vector indicated as a superimposed bar on each polar plot. (h) At the neural circuit level, the theta-oscillation-mediated neural synchrony in the VTA-mPFC circuit that emerged during punishment-free actions declined during punishment-prone actions. Neural synchrony mediated by the theta oscillation may subserve binding of the VTA-mPFC neurons responding to the appetitive action into the “appetitive” neural network. Our observation of decline in theta-mediated neural synchrony may reflect reduced activation of the appetitive neural network in the presence of punishment contingency.

In sum, we found a coherent theta oscillation temporarily synchronized the VTA-mPFC neural circuit during rewarded instrumental action. This synchrony declined as a function of punishment contingency (Figure 8h).

## Discussion

To unravel the VTA and mPFC neural representation of punishment contingent on goal-directed behavior, we engaged animals in an instrumental task where an action consistently procured a reward but probabilistically led to punishment. Simultaneous recording from VTA and mPFC demonstrated that these regions use multiple coding structures, involving spike-rate and LFP-mediated neural synchrony, to represent punishment contingent on goal-directed actions. VTA and mPFC single neurons encoded the same action differently if that action was punishment-free versus punishment-prone, suggesting that these neurons encode the contingency between action and punishment. At the network level, coherent theta oscillations that emerged in mPFC and VTA during punishment-free actions declined during punishment-prone actions, indicating that risk of punishment disrupts the synchrony between the two regions associated with an appetitive state.

### Neurons encode action-punishment contingency on different timescales

Using a novel behavioral paradigm, we modeled anxiety-like changes in instrumental actions as a function of action-punishment contingency. We found that the vast majority of VTA and mPFC neurons encode punishment by modulating their firing rates. Critically, the neuronal encoding was most pronounced at the time of the action as compared to other task events. This strongly suggested that neurons in both regions represented contingency between action and punishment.

Punishment influences behavior, and neuronal representation associated with that behavior, on multiple timescales (Cohen et al., 2015; Somerville et al., 2013). On short timescales, real-time neural processing of punishment may be important to signal contingency of punishment on a specific event in order to promote rapid behavioral adaptation. Our data suggest that VTA DA neuronal signaling may be involved in this context. DA neurons displayed phasic excitatory responses tightly linked to each of the task events (cue, action, and reward). Importantly, the DA neuronal encoding of punishment was concentrated around the time of the action compared with other task epochs, suggesting that DA neuronal signaling of punishment may primarily reflect the action-punishment contingency. On longer timescales, punishment can elicit persistent changes in motivational and emotional states – e.g., changes in mood. We found that mPFC and VTA non-DA neurons display temporally diffuse encoding of punishment within the peri-action window. Likewise, many of the mPFC and non-DA neurons showed significant modulation of their baseline firing rates, suggesting that these neurons may encode punishment with persistent changes in activity. This sustained change in activity may be responsible for longer-lasting affective impact of punishment.

### Neurons display bidirectional encoding of action-punishment contingency

Different subpopulations of punishment-encoding neurons in both regions encoded punishment by increasing or decreasing their peri-action firing rates. The direction of neuronal responses to appetitive vs aversive events is thought to carry information about motivational properties encoded by the neuronal activity. While heterogeneous response patterns have been widely observed in PFC neuronal encoding of punishment (Kobayashi et al., 2006; Matsumoto et al., 2007; Seo and Lee, 2009; Ye et al., 2016), there is some debate on whether the VTA DA neurons encode punishment with excitatory or inhibitory responses (Bromberg-Martin et al., 2010; Schultz, 2016). It has been demonstrated that reward prediction error (RPE)-coding DA neurons respond to appetitive (better-than-predicted) vs aversive (worse-than-predicted) events by excitation and inhibition, integrating information about appetitive and aversive events into a common currency of value (Eshel et al., 2016; Matsumoto et al., 2016; Mileykovskiy and Morales, 2011; Mirenowicz and Schultz, 1996; Roitman et al., 2008). We found that a subset of VTA DA neurons conforms to this pattern. These neurons responded to punishment-free (purely appetitive) actions with phasic excitation, which decreased as a function of punishment contingency.

In contrast, a greater proportion of DA neurons showed excitatory encoding of punishment; i.e., they treated appetitive and aversive components in the same direction, and responded to actions prone to punishment with further excitation. We observed that excitatory encoding of punishment contingency was more predominant among DA neurons, suggesting that the contingency of punishment is not simply encoded as reduced value of the action. Previous studies reported a subpopulation of DA neurons showing similar excitatory responses to appetitive and aversive stimuli (Brischoux et al., 2009; Joshua et al., 2008; Matsumoto and Hikosaka, 2009; Valenti et al., 2011). An excitatory DA neuronal encoding of aversive or neutral events has been interpreted as conveying motivational salience (Bromberg-Martin et al., 2010; Matsumoto and Hikosaka, 2009), detection (Nomoto et al., 2010), intensity (Fiorillo et al., 2013) of a sensory event, as well as generalization effect of rewarding stimuli (Kobayashi and Schultz,2014). Combination of these factors may comprise the excitatory DA neuronal encoding of punishment-prone actions we observed. Furthermore, the DA neuronal encoding of aversion has been suggested to depend on animals’ behavioral state. A recent study demonstrated that DA neurons encoded aversive events with inhibition in a low reward context but with biphasic responses in a high reward context (matsumoto et al., 2016). In addition, contrasting DA neural responses were observed based on how animals responded to an aversive event. DA concentration in the ventral striatum increased or decreased when rats displayed active avoidance or passive reaction (freezing) to shock-predicting cues (Oleson et al., 2012; Wenzel et al., 2015). These patterns of state dependency in DA neural responses to aversion are in line with our observation of excitatory encoding of punishment when risk was taken for reward in a highly rewarding context. Thus, our results are in support of the view that there are different modes of DA neuronal signaling of aversion depending on animals’ behavioral state and/or the reward availability in the environment.

Punishment is an event of negative valence associated with avoidance but of high salience deserving prioritized attention. Both aspects of punishment need to be represented for appropriate behavioral coping. Our observation of bidirectional encoding of punishment by distinct subpopulations of VTA DA, non-DA, and mPFC neurons suggests that different neuron groups may convey diverse motivational and emotional properties of punishment.

### VTA-mPFC neural synchrony declines with punishment

At the network level, we observed that coherent theta oscillation synchronized VTA and mPFC specifically during punishment-free actions, effectively phase-modulating the neuronal spike activity in the two regions. Analyses of the temporal relationship indicated that the neural synchrony arose in the VTA-to-mPFC direction. That is, the VTA driven bottom-up theta oscillation entrained mPFC LFPs and neuronal spike activity during punishment-free actions. The theta oscillation preferentially entrained DA neurons but much fewer non-DA neurons in the VTA. Considering the phasic excitatory responses of DA neurons during action, the theta oscillation-mediated neural synchrony may promote phase-coupling between VTA DA and mPFC neurons selectively during punishment-free actions. This phase-coupling diminished during punishment-prone actions. Thus, prediction of punishment can be inferred by dual alterations in the phase and the rate of the DA neurotransmission in the mPFC (model in Figure 8h).

Our data showing preferential DA (vs non-DA) neuronal phase-locking with VTA and mPFC theta oscillations supports the theoretical model suggesting that the VTA DA input may play a crucial role for cortical theta oscillations (Buhusi and Meck, 2005). In addition, our data is consistent with previous studies showing that inhibiting DA D1 receptors in mPFC diminishes theta oscillations (Parker et al., 2014). While somewhat out of the scope of the present study, it is noteworthy that theta oscillations in the mPFC and/or VTA have been related to the hippocampal theta oscillation (Hyman et al., 2005; Jones and Wilson, 2005; Siapas et al., 2005; Sirota et al., 2008). Local infusion of DA in mPFC enhances theta oscillations and coherence between mPFC and hippocampus (Benchenane et al., 2010), while both mPFC and VTA neurons exhibit phase coherence to the hippocampal theta oscillation (Fujisawa and Buzsaki, 2011). Thus, the theta-mediated neural synchrony we observed in the VTA-mPFC circuit may be coupled with the hippocampal theta oscillation.

Recent studies have reported slow (4 Hz) oscillation in mPFC during fear-conditioned freezing in synchrony with other regions including the amygdala (Dejean et al., 2016; Karalis et al., 2016b; Likhtik et al., 2014). Considering the distinct behavioral states associated with 4 Hz oscillations in these studies and theta oscillations observed in our study, these findings together suggest that the mPFC may be entrained by distinct bands of oscillations in appetitive vs aversive states. However, the 4 Hz oscillation in the VTA-mPFC circuit has also been associated with working memory (Fujisawa and Buzsaki, 2011). The fast vs slow mPFC oscillations occurring in appetitive vs aversive states may arise in preferential synchrony with VTA or amygdala in appetitive vs aversive states, thereby the bottom-up information transfer from the two subcortical regions can be routed depending on the behavioral context. This scenario would predict that the theta-oscillation-mediated mesoprefrontal synchrony would diminish in the presence of punishment, whereas a 4 Hz oscillation may arise in the mPFC. In accord with this, our results showed that mPFC theta-oscillation-mediated synchrony significantly declined as a function of action-punishment contingency. We did not observe the emergence of 4 Hz oscillation presumably because, unlike fear conditioning paradigms, our task involved instrumental actions. Entrainment of a neural circuit with varying frequency oscillations as a function of a task variable has been widely observed in sensory cortical circuits (Bosman et al., 2012; Jia et al., 2013). Such frequency modulation, along with the power modulation, could promote selection and binding of task-relevant neuronal ensembles to give rise to a functional neural network (Fries, 2015). Likewise, our data may reflect the rise and fall of coherent VTA and mPFC neuronal ensembles that may promote a flexible control of instrumental behavior as a function of punishment contingency.

The neural synchrony mediated by different bands of oscillations in distinct behavioral states may also implicate neuron groups in the other mesocorticolimbic structures such as nucleus accumbens, amygdala, hippocampus, and lateral habenula that are critical for appetitive and aversive behaviors. Mounting evidence suggests that distinct VTA and mPFC neuron groups within the same region selectively respond to appetitive vs aversive events (Lammel et al., 2012; Ye et al., 2016). Importantly, these studies have shown that the neuron groups differentially tuned for appetitive vs aversive events display discrepant patterns of input-output connectivity within the mesocorticolimbic system (Howe and Dombeck, 2016; Lammel et al., 2012; Parker et al., 2016; Roeper, 2013; Ye et al., 2016). This projection specificity may be the foundation for the selective recruitment of distinct neuron groups in distinct-valence experiences. The anatomical connectivity alone, however, may not be sufficient to bind the neuron groups tuned to appetitive or aversive events into a functional neural network in a timely manner. Therefore, the neural synchrony mediated by coherent oscillations, such as that demonstrated here, may play a key role for the rise and fall of the functional neural networks depending on the behavioral context.

In conclusion, proper encoding of punishment and its contingency on an action is fundamental to adaptive behavior and survival. Our data reveal dynamic coding schemes of the VTA-mPFC neural circuit in representing risk of punishment and punishment-based modulation of rewarded actions.

## Experimental procedure

### Subjects and surgical procedure

Male Long Evans rats weighing 300~400 g (Harlan) were singly housed on a 12 h light/dark cycle (lights on at 7 p.m.). All data were collected during the dark cycle. Microelectrode arrays were surgically implanted in ipsilateral mPFC and VTA (N = 10) or bilateral mPFC (N = 4) of isoflurane-anesthetized rats (Figure 2a). All mPFC electrode arrays were placed in the prelimbic subregion of the mPFC. The following coordinates relative to the bregma were used: mPFC = AP +3.0 mm, ML 0.7 mm, DV 4.0 mm; VTA = AP −5.3 mm, ML 0.8 mm, DV 8.2 mm (Paxinos and Watson, 1998). Behavioral training began after 1 week of postsurgical recovery. At the completion of all recordings, rats were anesthetized with 400 mg/kg chloral hydrate and perfused with saline and 10 % buffered formalin. Coronal brain slices of mPFC and VTA were collected and cresyl-violet stained. Placements of electrode arrays were verified under a light microscope. All procedures were in accordance with the National Institute of Health’s Guide to the Care and Use of Laboratory Animals, and were approved by the University of Pittsburgh Institutional Animal Care and Use Committee.

### An instrumental task with varying punishment-action contingency

After the postsurgical recovery, rats were kept at 85 % of their free-feeding weight on a restricted diet of 13 g food pellets a day with free access to water. In an operant chamber, rats were fully trained to make an instrumental nose poke to the cue port to receive a sugar pellet at the food trough located in the opposite side of the chamber on the fixed ratio schedule of one i.e., FR1 (Figure 1a-b). After completion of three FR1 sessions consisting of 150 trials in 60 mins, rats were trained with the task consisting of three blocks with varying degrees of action-punishment contingency (50 trials per block). Each block was assigned an action-punishment contingency of 0, 0.06, or 0.1 – i.e., the conditional probability of receiving an electrical foot shock (0.3 mA, 300 ms) given an action. The action–reward contingency was kept at 1 across all training and recording sessions; i.e., every nose poke procured a reward even in the shock trials. To minimize generalization of the action-punishment contingency across blocks, they were organized in an ascending shock probability order – Block1: 0, Block2: 0.06, Block3: 0.1, interleaved with 2-min timeout between blocks. In block 2 and 3 of each session, 3 and 5 trials were pseudo-randomly selected and followed by an electrical foot shock. No explicit cue was provided on shock trials to keep the shock occurrence unpredictable. The cue onset only signaled initiation of a trial. Animals were informed of the block shift by the 2-min darkened timeout in between blocks. In addition, the first shock trial of block 2 and the first two shock trials of block 3 were randomly selected from the initial 5 trials of each block. Also, animals completed two sessions of this task before the recording session, thus the shock occurrence and the task design including the ascending punishment contingency were not novel to them at the time of the recording session. All training and recording sessions were terminated if not completed in 180 mins, and data from the completed sessions only were analyzed. Animals displayed stable behavioral performance overall without any sign of contextual fear conditioning, since they performed fearless in the safe block across all sessions. In addition, there was no evidence for habituation to the shock as they showed equivalent punishment-based behavioral changes across sessions. For the diazepam pretreatment experiment, a separate group of rats (N = 9) were trained using abovementioned procedure, and they underwent three test sessions with intraperitoneal pretreatment of saline – diazepam (2 mg/kg, Hospira, Inc.) – saline. Injected animals were returned to their home cage for 10 minutes before they were placed in the operant chamber. Three days of washout period was allowed between sessions.

### Electrophysiology

Single-unit activity and local field potentials (LFPs) were recorded simultaneously using a pair of eight channel Teflon-insulated stainless steel 50 μm microwire arrays (NB Laboratories). Unity-gain junction field effect transistor headstages were attached to a headstage cable and a motorized commutator nonrestrictive to the animals’ movement. Signals were amplified via a multichannel amplifier (Plexon). Spikes were bandpass filtered between 220 Hz and 6 kHz, amplified ×500, and digitized at 40 kHz. Single-unit activity was then digitally high-pass filtered at 300 Hz and LFP were low-pass filtered at 125 Hz. Continuous single-unit and LFP signals were stored for offline analysis. Single units were sorted using the Offline Sorter software (Plexon). Only the single-units with a stable waveform throughout the recording session were further analyzed. If a unit presented a peak of activity at the time of the reference unit’s firing in the cross-correlogram, only either of the two was further analyzed.

### Neural data analysis

Single unit and LFP data analyses were conducted with Matlab (MathWorks) and SPSS statistical software (IBM). For single unit data analyses, 1-ms binned spike count matrix of the peri-cue, action, and reward periods (starting 2 s before each event and ending 2 s after each event) were produced per unit. The baseline period was a 2-s time window beginning 2.5 s before the cue onset. For all neural data analyses, the trials with shock delivery (three and five trials for block 2 and 3, respectively) were excluded as single-unit and LFP signals in these trials were affected by electrical artifacts during shock delivery.

*Trial-averaged firing-rate analysis.* Spike count matrices were further binned using a 200 ms rectangular moving window with steps of 50 ms within the −2 to 2 s epoch aligned to the task event occurring at time = 0 for the firing rate analysis. Binned spike counts were transformed to firing rates and averaged across trials. The trial-averaged firing rate of each unit was Z-score normalized using the mean and standard deviation of its baseline firing rate.

*VTA cell classification.* The VTA single units were classified into putative dopamine (DA) or non–dopamine (non-DA) neurons based on two criteria. First, units whose mean baseline firing rate slower than 12 Hz, waveform width greater than 1.2 ms were considered as potential DA units (Grace and Bunney, 1984; Kim et al., 2015; Schultz and Romo, 1987). This traditional classification, however, has been suggested to be potentially inaccurate (Margolis et al., 2006). Thus, the second criterion utilized the neuronal reward response properties for the putative DA and non-DA cell identification. Receiver-operating characteristic (ROC) curves were calculated by comparing the distribution of firing rates across trials in 100 ms bins (starting 0.5 s before reward delivery and ending 1 s after reward delivery) to the distribution of baseline firing rates. Principal component analysis was conducted using the singular value decomposition of the area under the ROC (auROC). Units were mapped in the 3-d space comprising the top three principal components. Within the 3-d PC space, unsupervised clustering was conducted by fitting Gaussian mixture models using the expectation-maximization algorithm. This method found two clusters: one with phasic excitation to reward (Type 1), one with sustained excitation or suppression to reward (Type 2) (Figure S1c-e). Units in the former class were classified as putative DA units, as previous studies have shown that optogenetically tagged dopamine neurons displayed similar phasic excitatory reward responses (Cohen et al., 2012; Eshel et al.,2015). Taken together, we defined a VTA unit satisfying both criteria as a putative DA unit and a unit that met either or none of the criteria as a putative non-DA unit (Figure S1). mPFC units were not classified based on their firing and spike-waveform properties. Only 2 out of the total 293 mPFC single units had the mean baseline firing rates higher than 20 Hz, thus few fast-spiking interneurons should be included in our data analysis.

*Spike rate selectivity.* To quantify single neuronal encoding of blockwise action-punishment contingency, we computed a bias-corrected percent explained variance (ωPEV) statistic with binned spike counts calculated in a 200 ms rectangular window moving with steps of 50 ms within the 2-s peri-event epochs (-1 to 1 s with an event occurring at time = 0).

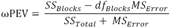

where SS_Blocks_ and SS_Total_ are the between-blocks (action-punishment contingency) and total sums of squares, df_Blocks_ is degrees of freedom, and MS_Error_ is the mean squared error. This formulation resulted in an unbiased metric with an expected value of zero when there is no difference across blocks (Buschman et al., 2012; Keren, 1979). A unit was determined to encode action-punishment contingency if its peri-event ωPEV surpassed ‘the global ωPEV band’, which was defined as the upper bound of the 99 % confidence interval of the trial-shuffled (1,000 times) surrogate ωPEV distribution – i.e., fewer than 1 % of the trial-shuffled ωPEVs crossed the global band across all time bins in the peri-event epoch (α = 0.01). To find the global ωPEV band, we computed the mean and standard deviation of the trial-shuffled ωPEV distribution. By stepping up from the mean by one-hundredth of the standard deviation, we found the pointwise band at each time bin and the global band across time bins both at α = 0.01 (Figure S2). This approach effectively resolves the issue of multiple comparisons that can arise as statistical comparisons made separately across multiple time bins increase the rate of false rejection of the null hypothesis (Fujisawa et al., 2008). We repeated this analysis using the mutual information metric, and found that the two metrics yielded similar results.

*Linear regression analysis.* For a standardized quantification of the individual neuronal encoding of action-punishment contingency in peri-event epochs, we computed the standardized regression coefficient of the following linear regression model for each unit:

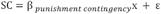

where SC denotes binned spike counts calculated in a 200 ms moving window with steps of 50 ms, *β_punishment contingency_* regression coefficients for the independent variable, blockwise punishment contingency (1, shock prob. = 0; 2, shock prob. = 0.06; 3, shock prob. = 0.1), respectively. The regression coefficients were standardized by β×(S_x_/ S_y_), where *S_x_ S_y_* denote the standard deviations of independent and dependent variables, respectively.

*LFP power spectra and coherence.* The local field potential (LFP) power spectral densities were quantified using the chronux routine mtspecgramc (Bokil et al., 2010). Briefly, the LFP time series within the peri-event epochs were Fourier transformed in a 500 ms moving window with steps of 50 ms with the multi-taper method applied:

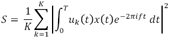

where u_k_(t) is the multi-tapers (9 tapers were used), x(t) is the LFP time series in a moving window. The baseline normalized power spectra (Z-score) were calculated using the mean and standard deviation of the baseline power spectra across trials. In addition, we inspected the trial-by-trial spectro-temporal representations of LFP time series applying the continuous wavelet transform. We confirmed that comparable representations were attained by the Fourier-and wavelet-based time-frequency analyses.

The magnitude squared coherence (MSC) between time series recorded from mPFC and VTA was calculated in the same moving window with the 9 multi-tapers applied using the chronux routine cohgramc. Briefly, the MSC was quantified as:

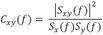

where S_xy_(f) is the cross spectral density of LFP time series in the two regions, and *s_x_(f),S_y_(f)* are the autospectral density for each region.

*Bivariate Granger causality analysis.* To examine mutual influences (directionality) between LFP oscillations in the two regions, we quantified Granger causality between the simultaneously recorded peri-action LFP traces (-2 to 2 s around the action occurring at time = 0). The bivariate Granger causality (G-causality) infers causality between two time series data based on temporal precedence and predictability (Barnett and Seth, 2014; Granger, 1969). That is, a variable X_1_ ‘Granger causes’ a variable *X_2_* if information in the past of X_1_ helps predict the future of *X_2_* with better accuracy than is possible when considering only information already in the past of X itself. In this framework, two time series *Xi(t)* and *X_2_(t)* recorded from mPFC and VTA can be described by a bivariate autoregressive model:

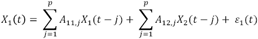

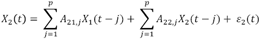

where *p* i s the model order (the maximum number of time-lagged observations included in the model), which was estimated by the Akaike information criterion. We then estimated parameters of the model; A contains the coefficients of the model, and ε_1_ ε_2_ are residuals (prediction errors) with covariance matrix Σ for each time series.

Once the model coefficients *A*_j_and Σ are estimated, the spectral matrix can be obtainedas:

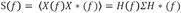

Where the asterisk denotes matrix transposition and complex conjugation, Σ is the noise covariance matrix, and the transfer function H(f) is defined as the inverse matrix of the Fourier transform of the regression coefficients:

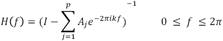

The spectral G-causality from 1 to 2 is then obtained by:

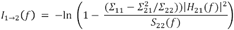

The spectral G-causality measure lacks known statistical distribution, thus a random permutation method was used to generate a surrogate distribution, by which the upper bound of the confidence interval was found at α = 0.001. This procedure was implemented using an open-source matlab toolbox, the Multivariate Granger Causality Toolbox (Barnett and Seth, 2014).

*LFP-Spike phase-locking analysis.* To quantify the individual neuronal spike time synchrony with the local and interregional theta oscillations were quantified as follows. The mPFC and VTA LFPs during the baseline and peri-action 4-s time windows were bandpass filtered to isolate oscillations within 5 – 15 Hz frequency range. The instantaneous phase of each filtered LFP segment was determined using the Hilbert transform and each spike was assigned the phase of its contemporaneous LFP segment. The phase-locking value (PLV) of each unit was defined as the circular concentration of the resulting phase angle distribution, which was quantified as the mean resultant length (MRL) of the phase angles. The MRL is the modulus of the sum of unit vectors indicating instantaneous phases of each spike occurrence normalized by the number of spikes, thus the MRLs were bound to take a value between 0 (no phase-locking) to 1 (perfect phase-locking).

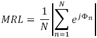

Where Φ_n_ represents the phase assigned to n^th^ spike occurrence, and *N* is the total number of spikes. Since the MRL statistic is sensitive to the number of spikes, we calculated MRL 1,000 times with 100 spikes of each unit randomly selected for each iteration, and the PLV was the MRL averaged across all iterations. As comparing PLVs across blocks with varying action-punishment contingency was of our central interest, PLV was computed per each block. Only the units with their peri-action spike counts within each block greater than 100 in all three blocks were included in the phase-locking analysis. Units passing the Rayleigh’s z-test at α = 0.05 were determined to be significantly phase-locked. The directionality of the LFP and spike phase relationship was inferred by a time-lagged phase-locking analysis, in which the spike times were shifted relative to the LFP time series, stepping by 4 ms within the range of −100 to 100 ms (Likhtik et al., 2014; Spellman et al., 2015). At each time lag, the PLV of each single unit and its significance were assessed, and the maximum PLV across all time lags was found for each unit. We repeated the analysis with different time lags and analysis windows, and confirmed that the results were very similar across different parameters.

### Statistical analysis

Parametric statistical tests were used for z-score normalized data and non-normalized data that are conventionally tested using a parametric test. Nonparametric approaches, such as conventional nonparametric tests or bootstrapping were used for a hypothesis test of data, of which statistical distribution is unknown, e.g. phase-locking values (PLVs). For all tests, the Greenhouse-Geisser correction was applied as necessary due to violations of sphericity. All statistical tests were specified as two-sided. Multiple testing correction was applied for all tests including multiple comparisons using the Bonferroni correction.

---

**Supplementary Figure 1.**
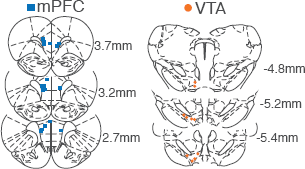
Histologically verified placements of mPFC and VTA electrodes. We recorded activity of ipsilateral VTA and mPFC (N=10) or bilateral mPFC (N=4). Coordinates are relative to the bregma.

**Supplementary Figure 2.**
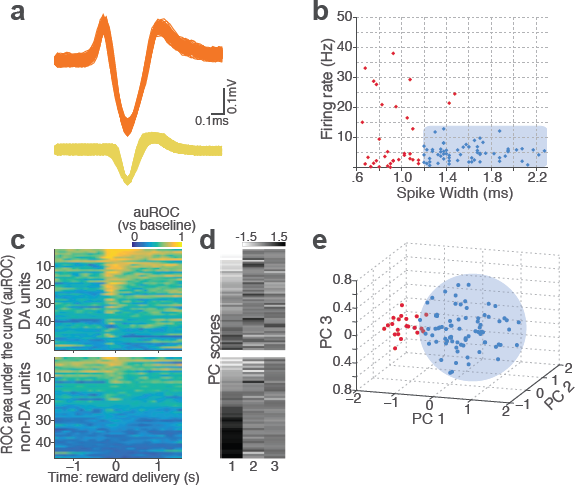
Classification of VTA single units to putative DA or putative non-DA units. (a) Representative spike waveforms of a putative DA (top) and a non-DA (bottom) units.(b) Units were first classified based on their mean baseline firing rate and width of the spike waveform. Units whose mean baseline firing rate slower than 12 Hz, waveform width greater than 1.2 ms were considered to be putative DA units (blue circles). (c) To characterize each unit’s reward response, ROC curves were calculated by comparing the firing-rate distributions of reward delivery vs baseline epochs. (d) Principal component analysis (PCA) was conducted on auROC values. (e) Units were mapped onto a 3-d space comprising the top three principal components. Unsupervised clustering was conducted by fitting Gaussian mixture models which yielded two clusters of units: one with phasic excitation to reward (blue circles), the other with sustained excitation or suppression to reward (red circles). Units in the former cluster were classified as putative DA units. Only the units satisfying both criteria (b) and (e) were finally labelled as putative DA units, and the rest of units were putative non-DA units.

**Supplementary Figure 3.**
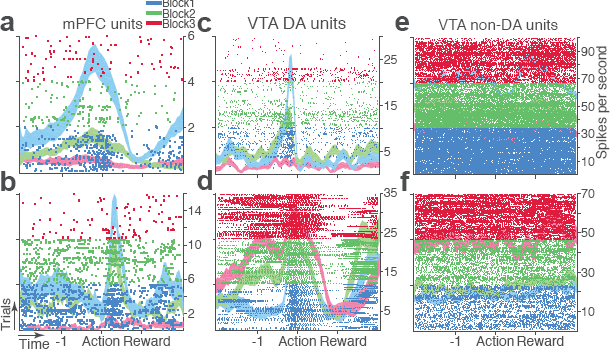
Representative punishment-encoding mPFC (a-b), VTA DA (c-d), and non-DA (e-f) single units. In each plot, data are for 150 trials of action with three different levels of punishment contingency. Each row represents each trial. Ticks mark spike times. The horizontal axis represents time around the action occurring at time = 0. Spike density functions of different blocks are superimposed as mean ± s.e.m. (shaded area).

**Supplementary Figure 4.**
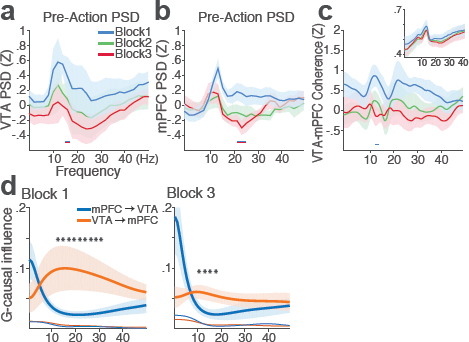
mPFC and VTA theta oscillations did not change across blocks in the absence of punishment. (a) Mean ± s.e.m. (shaded area) normalized VTA PSDs of each block corresponding to 1 s pre-action epoch. Dual-colored bars below indicate significant pairwise differences at corresponding frequency bins according to post hoc analyses (p < 0.05). (b) Normalized mPFC PSDs. (c) Normalized VTA-mPFC LFP coherence of each block in the pre–action epoch. Insets represent non-normalized LFP coherences. (d) Granger-causality, representing mutual influences (directionality) between VTA and mPFC peri-action LFP time series in block 1 (left) & 3 (right). Blue and orange curves represent mPFC-to-VTA and VTA-to-mPFC Granger-causal influences, respectively. Shaded areas indicate s.e.m. Thin colored-lines below indicate upper bounds of confidence intervals (α = 0.001) acquired by the random permutation resampling of time bins. An asterisk indicates significant difference between bidirectional Granger-causal influences at the corresponding frequency bin (p < 0.05).

**Supplementary Figure 5.**
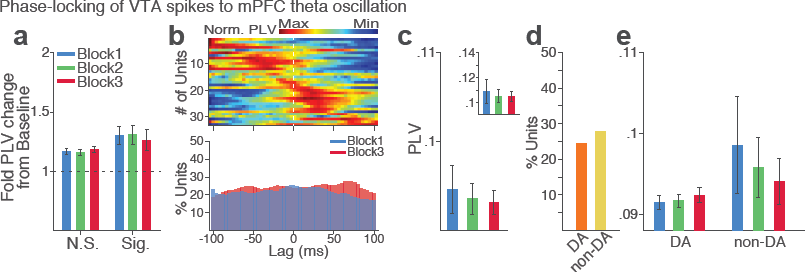
VTA single units show weak phase synchrony to the mPFC theta oscillation. (a) Fold change from baseline in the strength of the neuronal phase-locking. (b) Top, Normalized PLVs in block 1 for all phase-locked VTA units. The phase-locked units displayed their peak PLVs with negative or positive time lags, indicating weak phase modulation of VTA spike activity by the mPFC theta oscillation (Signed-rank test, *p* = 0.129). Bottom, Percentage of significantly phase-locked VTA units did not differ across blocks. (c) Mean ± s.e.m. PLVs across different blocks. No significant change was found across blocks (Signed-rank test, *p* values >0.33). (d) Percentage of phase-locked VTA DA vs non-DA units. (e) PLVs of VTA DA and non-DA units plotted separately. PLVs did not differ across blocks in both VTA cell types (Signed-rank test, *p* values > 0.25).

**Supplementary Figure 6.**
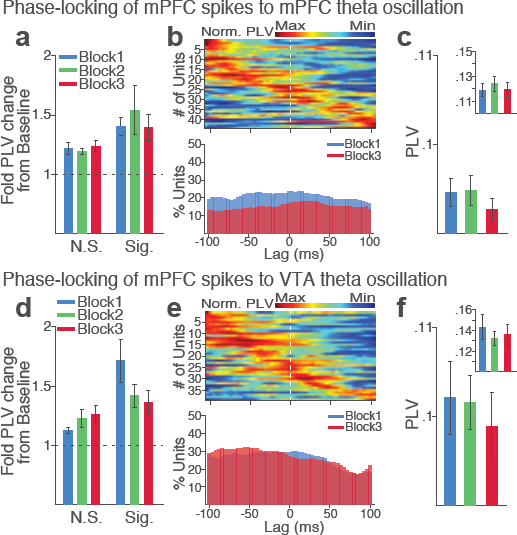
mPFC neuronal synchrony to mPFC and VTA theta oscillations did not change across blocks in the absence of punishment (No-shock control). (a-c) mPFC neuronal phase-locking to mPFC theta oscillation. (a) Fold change from baseline in the strength of the neuronal phase-locking during peri-action epoch in units that passed Rayleigh z-test (Sig.) and rest of the units (N.S.). (b) Top, Normalized PLVs in block 1 across a range of time lags for all phase-locked mPFC units, aligned by peak lags. Bottom, Percentage of significantly phase-locked mPFC units in block 1 vs 3 across a range of lags. (c) Mean ± s.e.m. PLVs across different blocks. Inset, PLVs including significantly phase-locked units only. The PLVs did not significantly differ across blocks (Signed-rank test, *p* values > 0.105). (d-f) mPFC neuronal phase-locking to the VTA theta oscillation. The PLVs did not significantly differ across blocks (Signed-rank test, *p* values ≥ 0.392).

**Supplementary Figure 7.**
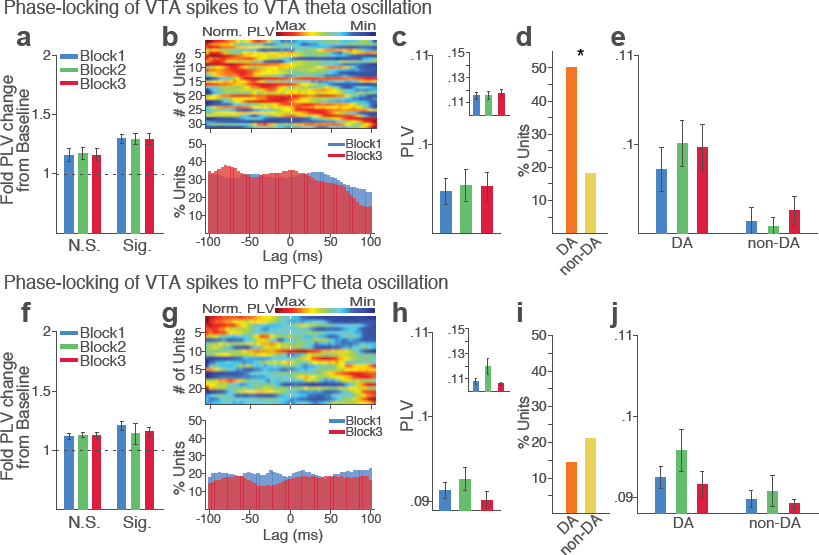
VTA neuronal synchrony to VTA and mPFC theta oscillations did not change across blocks in the absence of punishment (No-shock control). (a-e) VTA neuronal phase-locking to the VTA theta oscillation. (a) Fold change from baseline in the strength of the neuronal phase-locking during peri-action epoch in units that passed Rayleigh z-test (Sig.) and rest of the units (N.S.). (b) Top, Normalized PLVs in block 1 across a range of time lags for all phase-locked VTA units, aligned by s. Bottom, Percentage of significantly phase-locked VTA units in block 1 vs 3 across a range of lags. (c) Mean ± s.e.m. PLVs across different blocks. The PLVs did not significantly differ across blocks (Signed-rank test, *p* values > 0.743). Inset, PLVs including significantly phase-locked units only. (d) Percentage of phase-locked VTA DA vs non-DA units. Greater fraction of DA units (50 %) appeared to be phase-locked to the VTA theta oscillation compared with non-DA units (18 %) (Chi-square test, = 6.959, *p* = 0.008). (e)PLVs of VTA DA and non-DA units plotted separately. (f-j) VTA neuronal phase-locking to the mPFC theta oscillation. (h) The VTA neuronal phase-locking to the mPFC theta oscillation did not differ across blocks (Signed-rank test, *p* values > 0.355). Conventions are same as above.

## Author contributions

J.P. and B.M. designed research; J.P. performed experiments and analyzed data; J.P. and B.M. wrote the paper.

